# Mechanical evolution of metastatic cancer cells in three-dimensional microenvironment

**DOI:** 10.1101/2024.06.27.601015

**Authors:** Karlin Hilai, Daniil Grubich, Marcus Akrawi, Hui Zhu, Razanne Zaghloul, Chenjun Shi, Man Do, Dongxiao Zhu, Jitao Zhang

## Abstract

Cellular biomechanics plays critical roles in cancer metastasis and tumor progression. Existing studies on cancer cell biomechanics are mostly conducted in flat 2D conditions, where cells’ behavior can differ considerably from those in 3D physiological environments. Despite great advances in developing 3D *in vitro* models, probing cellular elasticity in 3D conditions remains a major challenge for existing technologies. In this work, we utilize optical Brillouin microscopy to longitudinally acquire mechanical images of growing cancerous spheroids over the period of eight days. The dense mechanical mapping from Brillouin microscopy enables us to extract spatially resolved and temporally evolving mechanical features that were previously inaccessible. Using an established machine learning algorithm, we demonstrate that incorporating these extracted mechanical features significantly improves the classification accuracy of cancer cells, from 74% to 95%. Building on this finding, we have developed a deep learning pipeline capable of accurately differentiating cancerous spheroids from normal ones solely using Brillouin images, suggesting the mechanical features of cancer cells could potentially serve as a new biomarker in cancer classification and detection.

## 1. Introduction

Biomechanical abnormalities in cells have been observed across various types of cancers during tumor development.^[1-3]^ Notably, profound studies have suggested that metastatic cancer cells are usually softer than normal cells.^[4-8]^ A reasonable explanation for this softening stems from the physical aspects of the metastatic process.^[9, 10]^ During metastasis, cancer cells have to migrate through highly confined regions such as dense extracellular matrices, surrounding tumor tissue, and the basement membrane of blood vessels; therefore, a deformable cell body facilitates the success rate of this process.^[11, 12]^ Despite the substantial progress in understanding the biomechanical regulation of cancer cells, a major limitation of existing studies is that the experiments were mostly conducted with cells grown on flat 2D environments or within rigid microfluidic devices, which intrinsically differ from 3D physiological environments thus could alter cancer cells’ behavior considerably.^[13-16]^ Therefore, to enhance the physiological relevance of the observations, it is essential to investigate the biomechanical phenotype of cancer cells within 3D microenvironments. Great advances have been made in developing physiologically relevant 3D *in vitro* models, such as spheroids and organoids, for cancer research.^[17-20]^ However, probing the mechanical properties of cancer cells in 3D conditions remains a significant challenge for conventional technologies, leaving the knowledge of this field largely unexplored.

Most of existing mechanical testing technologies, including atomic force microscopy or microcantilever based indentations, micropipette aspiration, microplate stretcher, and substrate stretcher, require physical contact thus are not directly accessible to cells in 3D conditions.^[21-23]^ Non-contact method such as optical tweezer has been successfully employed to probe the stiffness of single cancer cell in 3D culture^[24]^ and multicellular spheroids.^[25]^ However, the requirement for bead injection renders this method invasive and limits the probing to only a few points within a sample. In addition, mechano-microscopy based on optical coherence elastography has acquired elastic maps of spheroids in different environments with a lateral resolution of 5 μm and an axial resolution of 15 μm.^[26]^ Recently, an emerging non-contact technology, known as optical Brillouin microscopy, provides a promising solution to the current technological challenge.^[27-29]^ Brillouin microscopy is based on the physical principle of spontaneous Brillouin light scattering, where the scattered light undergoes a frequency shift, known as Brillouin shift, due to the localized interaction of the incoming light with the material.^[30]^ Since Brillouin shift is related to the elastic longitudinal modulus of the material, the optical readout of the frequency shift using a Brillouin microscope allows direct quantification of material’s mechanical properties. Consequently, a confocal Brillouin microscope can map material’s elasticity with subcellular resolution in a noncontact, noninvasive, and label-free manner.^[31-36]^ In the past decade, the feasibility of applying Brillouin microscopy to multicellular spheroids has been independently validated by several groups including us, demonstrating that this technology can conduct dense mechanical mapping in 2D/3D with high spatial resolution and high mechanical sensitivity.^[24, 37-41]^

In this work, to reveal the mechanical evolution of metastatic breast cancer cells during the early growth of tumor, we acquired mechanical images of cancerous spheroids using Brillouin microscopy and tracked their mechanical changes over eight days. We observed that, compared to their normal counterparts, cancer cells had distinct mechanical characteristics during sustained growth in the 3D microenvironment. Specifically, the average Brillouin shift of cancerous spheroids initially increased from Day 2 to Day 5, followed by a decrease by Day 8. In contrast, the Brillouin shift of normal spheroids continuously increased and were significantly higher than that of cancerous spheroids across all days. Furthermore, thanks to the dense mechanical mapping, we observed that cancerous spheroids exhibited unique mechanical distribution and intratumor heterogeneity. Using a well-established machine learning algorithm, we demonstrated that incorporating these biomechanical features with morphological features extracted from Brillouin images could significantly increase the classification accuracy of cancer cells from 74% to 95%. Building on this encouraging result, we developed a deep learning pipeline for cancer cell classification. We demonstrated that the model could accurately differentiate cancerous spheroids from their normal counterparts solely using Brillouin images. Collectively, these results shed light on developing new biomarkers based on Brillouin-derived biomechanical features for the detection of breast cancer.

## 2. Results

### 2.1 Longitudinal mechanical imaging of cellular spheroids using Brillouin microscopy

Tumor spheroids of the metastatic breast cancer cell line MCF10CA1h (M3) were grown in 3D culture based on established protocols.^[42, 43]^ The M3 cell line is well characterized as fully malignant:^[44]^ with cells demonstrating a 100% likelihood of metastasizing and forming tumors in mice following intravenous injection. For comparison, normal spheroids from the non-tumorigenic cell line MCF10A (M1), which shares a common genetic background, were cultured in a similar manner (**Figure 1A**). The mechanical properties of the spheroids were probed using a confocal Brillouin microscope. This microscope can acquire 2D/3D Brillouin shift images with subcellular resolution by scanning the sample with a focused laser beam (**Figure 1B-1D**, also see Experimental Section).

**Figure 1.**
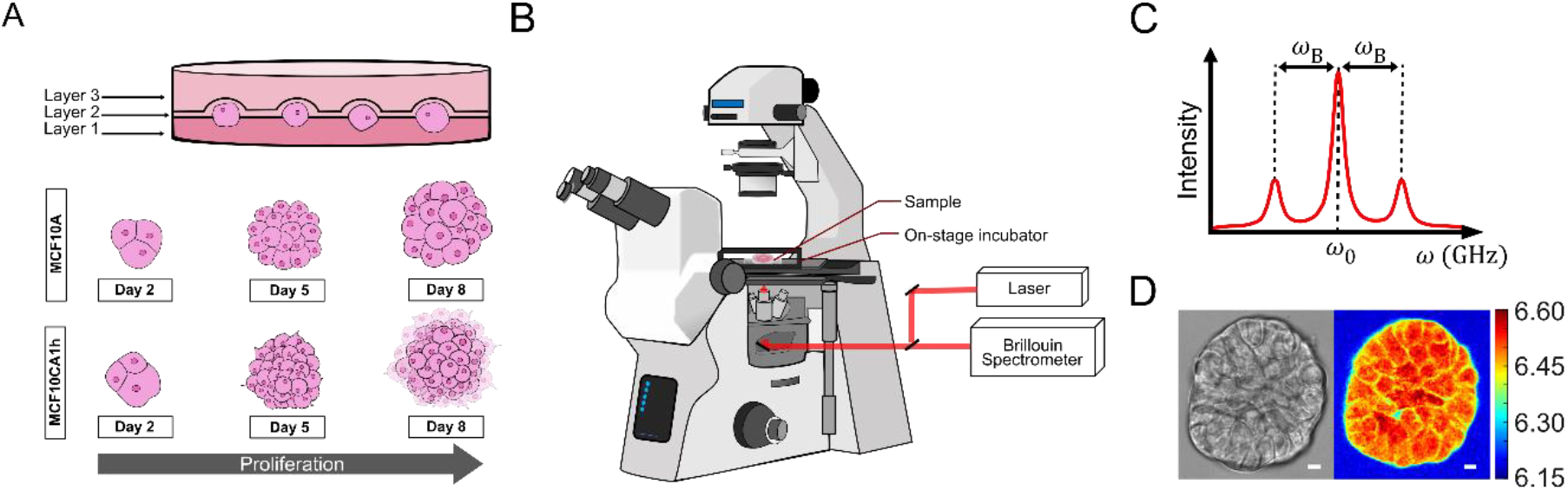
Biomechanical imaging of cellular spheroids using confocal Brillouin microscopy. (A) 3D on-top assay for the culture of spheroids over eight days. Layer 1 consists of 200 μl of Matrigel; layer 2 contains 500 μl assay medium with cell suspension; layer 3 contains 500 μl assay medium with 10% Matrigel. (B) Schematic of the self-built confocal Brillouin microscope. (C) Representative Brillouin spectrum, where *ω*_0_ is the frequency of the laser source and *ω*_*B*_ is Brillouin frequency shift measured by the Brillouin spectrometer. (D) Representative Brillouin image of a spheroid. The color bar represents the Brillouin frequency shift with a unit of GHz. Scale bar: 10 μm.

To track the mechanical changes of the spheroids during growth, Brillouin images of the same set of spheroids were acquired on Day 2, 5, and 8. The representative bright-field and Brillouin images of normal and cancerous spheroids are shown in Figure 2A. We first quantified the projection areas of the spheroids using the obtained Brillouin images (**Figure 2B**). Although we observed a rapid increase in size from Day 2 to Day 8 for both normal and cancerous spheroids, the difference in growth between the two types was not statistically significant. Furthermore, we tested the viability and the proliferation rate of the spheroids on Day 9, confirming that Brillouin microscopy is non-perturbative and does not cause photodamage (**Figure 2C-2D**).

**Figure 2.**
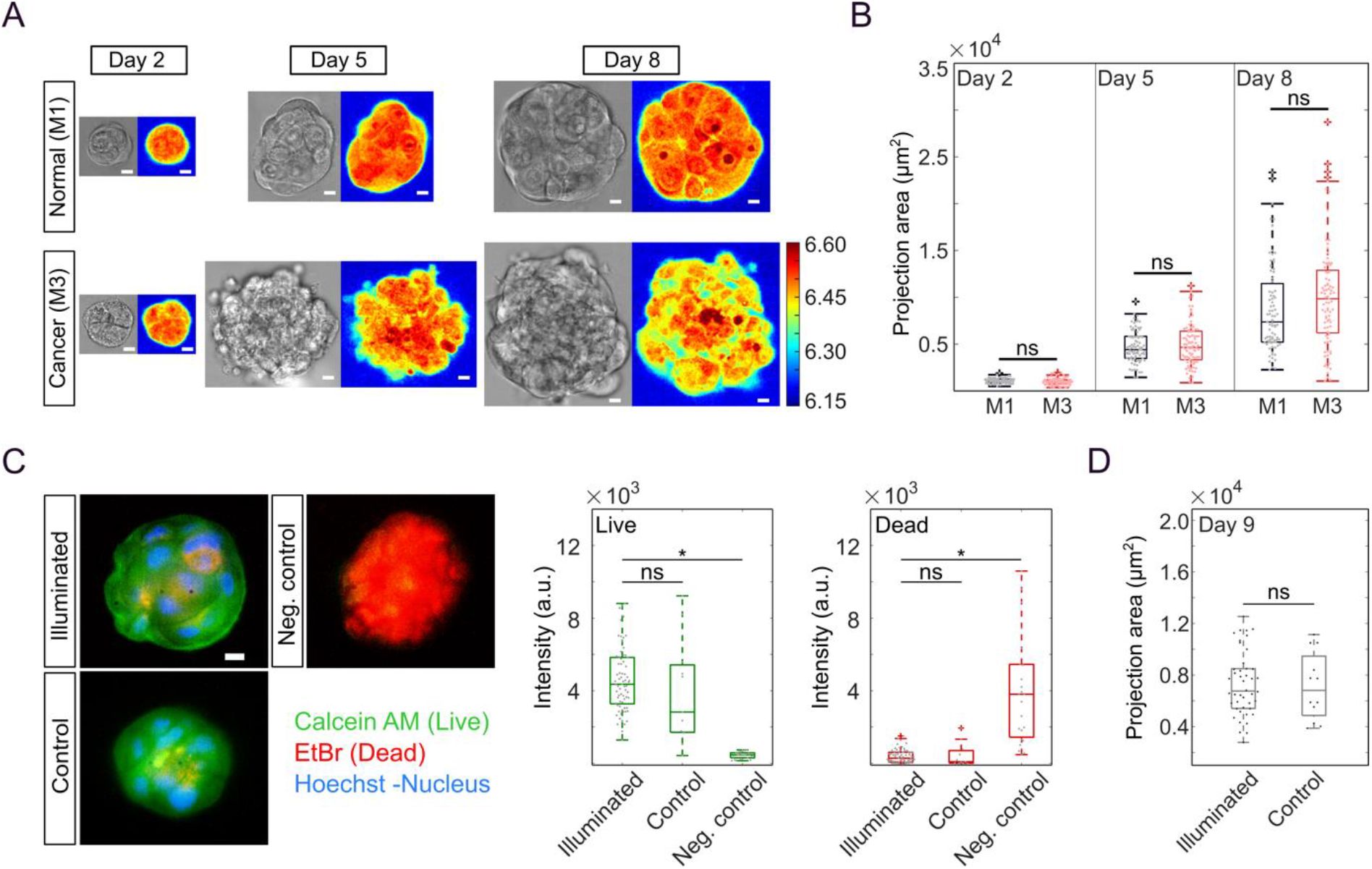
Longitudinal biomechanical Brillouin imaging of normal and cancerous spheroids. (A) Brillouin and bright-field images of a normal spheroid (M1) and a cancerous spheroid (M3) acquired on Day 2, 5, and 8. Scale bar: 10μm. (B) Projection area of normal (n=83) and cancerous spheroids (n=85). (C) Viability assay. Illuminated group, n=73; control group, n=11; negative control group, n=23. Scale bar: 20 μm. (D) Comparison of the projection area between illuminated (n=46) and control group (not illuminated by laser beam, n=12) on Day 9. \**p*<0.0001. n.s.: no statistical significance.

### 2.2 Cancerous spheroids showed distinct mechanical features during eight-day’s growth

In the case of normal spheroids (M1), we observed a consistent and significant increase in the average Brillouin shift from Day 2 to Day 8 (**Figure 3A**; **Figure S1**, Supporting Information). In contrast, for cancerous spheroids (M3), the average Brillouin shift initially increased significantly from Day 2 to Day 5, followed by a decrease from Day 5 to Day 8. As a result, the shift of cancerous spheroids on Day 8 showed no significant difference from that on Day 2. Notably, in all three days, normal spheroids had higher shift than their cancer counterparts, which is consistent with the observation on single cells (**Figure S2**, Supporting Information). To understand the spatial distribution of the intratumor elasticity, we segmented the spheroid into two zones: the core (region within 40% of the spheroid’s radius from the center) and the periphery (region outside of the core). We then quantified the average Brillouin shift of the two zones (**Figure 3B**). In all three days, we observed that the core had a consistently higher shift than the periphery for both cell lines, revealing a pronounced mechanical gradient from the spheroid’s center to its edge. This is consistent with previous observations.^[25, 39, 41]^ Furthermore, the changes in Brillouin shift over the course of spheroid growth show similar trends for both regions, suggesting that the global mechanical evolution is a common feature in the early-stage growth of the normal and cancerous spheroids.

**Figure 3.**
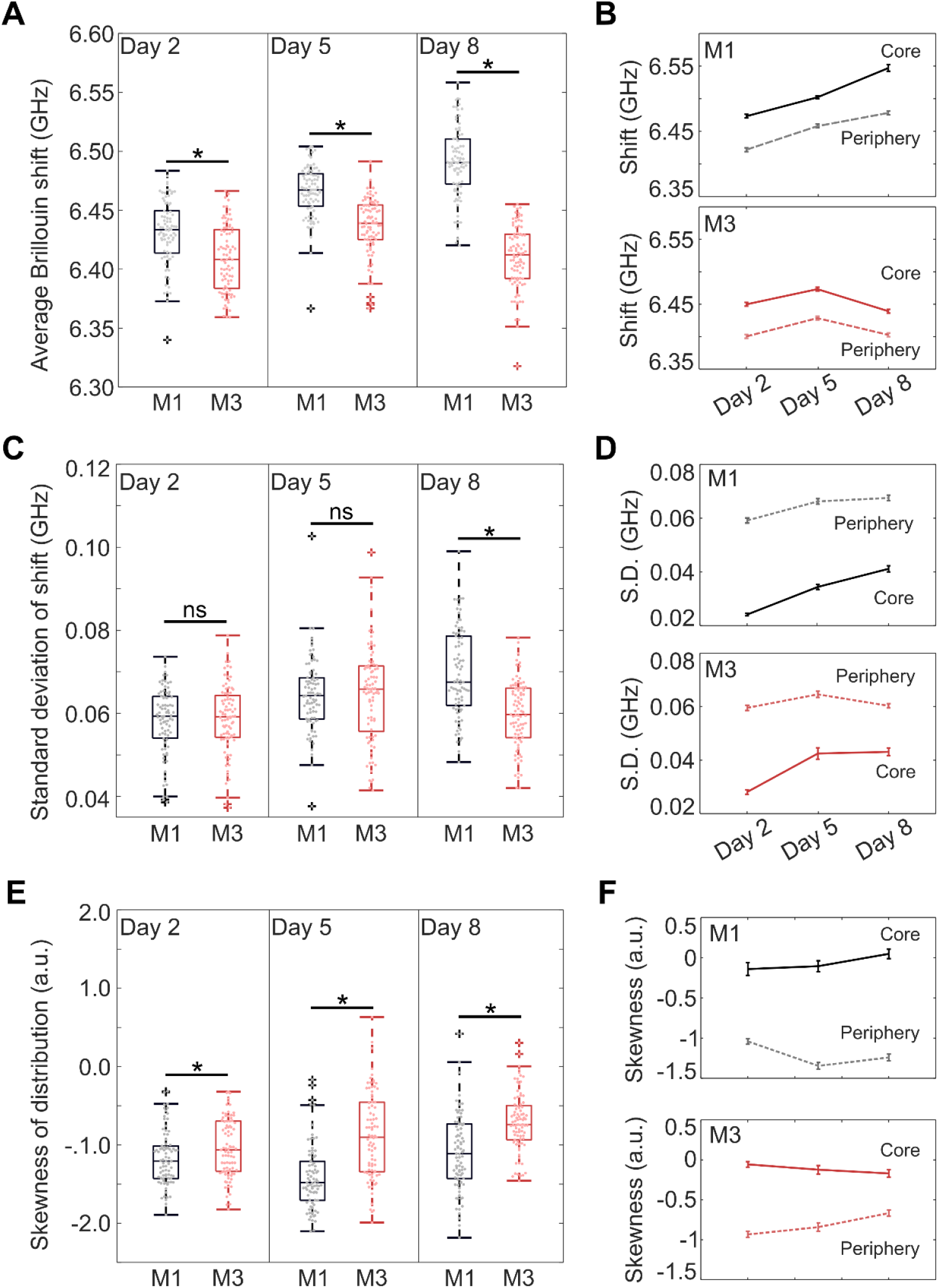
Mechanical evolution of normal and cancerous spheroids. (A) Average Brillouin shifts of normal (n=83) and cancerous (n=85) spheroids. \**p*-value for Day 2, 5, and 8 is 1.3×10^−6^, 2.9×10^−13^ and 5×10^−42^, respectively. (B) Average Brillouin shifts of the core and periphery regions of the spheroids. (C) Standard deviations of Brillouin shifts of normal and cancerous spheroids. \**p*=1.2×10^−10^. (D) Standard deviation of Brillouin shifts of the core and periphery regions. (E) Skewness of the histogram distribution. \**p*-value for Day 2, 5, and 8 is 0.02, 2.4×10^−11^, and 1.4×10^−7^, respectively. (F) Skewness of the core and periphery regions. Error bar represents standard error of the mean. n.s.: no statistical significance.

To further understand the biomechanical heterogeneity within the spheroid’s body, we quantified the standard deviation (SD) of Brillouin shifts for every spheroid (**Figure 3C**). Similar to the trends observed in the average shift, we noticed a continual increase in the SD for normal spheroids throughout the eight-day culture period, contrasted with an initial increase followed by a decrease in the SD for cancerous spheroids. However, the difference in the SD of the shift between two cell lines became significant only on Day 8, at what time normal spheroids exhibited greater SD of the shift compared to cancerous spheroids. This difference in mechanical heterogeneity on Day 8 is likely because normal cells start to form well-organized, acini-like structures, whereas cancer cells tend to migrate and form aggregations with varying degrees of disorganization.^[17, 25]^ Interestingly, we found that the periphery has a significantly higher SD of the shift compared to the core in both cell lines (**Figure 3D**). This suggests that the mechanical heterogeneity of the spheroids predominantly arises from the biomechanical organization at their periphery instead of their core.

To investigate the statistical distribution of the intratumor elasticity, we first plotted Brillouin shifts of all pixels within a spheroid into a histogram distribution and then measured its asymmetry (**Figure 3E&3F**). Here, the asymmetry was quantified by the parameter known as the skewness, which assumes a positive value when the distribution is left-tailed and a negative value when the distribution is right-tailed. For a distribution that is perfectly symmetrical, the skewness is zero. Briefly, we observed that all spheroids have negative skewness, suggesting that the distribution of shifts is skewed towards lower values, with the majority of pixels likely having shifts higher than the average value. For cancerous spheroids, we observed a consistent increase in the negative skewness from Day 2 to Day 8. However, normal spheroids showed a different trend, where the skewness first dropped by Day 5 and subsequently rebounded to a level comparable to that of Day 2 by Day 8. Intriguingly, we found that the asymmetric distribution of Brillouin shifts within a spheroid is primarily localized to the periphery, whereas the core showed a nearly symmetric distribution with the skewness close to zero. Importantly, across all three days, we observed that cancerous spheroids exhibited consistently less asymmetry in the mechanical distribution compared to normal spheroids. The unique skewness features of cancerous versus normal spheroids could potentially serve as an independent metric, distinct from the average and the SD of the shift, for differentiating between the two cell populations.

### 2.3 Mechanical phenotype serves as a new biomarker for cancer cell classification

Both the morphology and the physical properties of cells are associated with their physiological status and function. Previous studies have demonstrated that cellular morphology is a promising method for predicting the metastatic potential of cancer cells.^[45, 46]^ Here, we investigated the potential of Brillouin-derived mechanical features to enhance the accuracy of cancer cell classification when incorporating with the morphological features. To achieve this, we first extracted five morphological features and three mechanical features from the Brillouin image of every spheroid (**Table 1**). Next, we classified cancer and normal spheroids of each day by employing a well-established machine learning algorithm named support vector machine (SVM).^[47]^ We found that relying solely on morphological features yields a prediction accuracy of 65% on Day 2, which increased to 74% by Day 8 (**Figure 4**). Remarkably, by integrating both morphological and mechanical features, we observed a substantial improvement in prediction accuracy across all three days. Specifically, the classification accuracy on Day 8 surged to as high as 95%. In short, our study demonstrates the significant potential of Brillouin-derived mechanical features as new biomarkers for the classification of cancer cells.

**Table 1.** Morphological and mechanical features extracted from Brillouin images.

| Morphological features |  |  |  |  | Mechanical features |  |  |
| --- | --- | --- | --- | --- | --- | --- | --- |
| Area | Circularity | Solidity | Aspect ratio | C.V. of radii of convex hull <sup>a)</sup> | Average Brillouin shift | S.D. of Brillouin shift <sup>b)</sup> | Skewness |

<sup>a)</sup> C.V.: coefficient of variation; <sup>b)</sup> S.D.: standard deviation.

**Figure 4.**
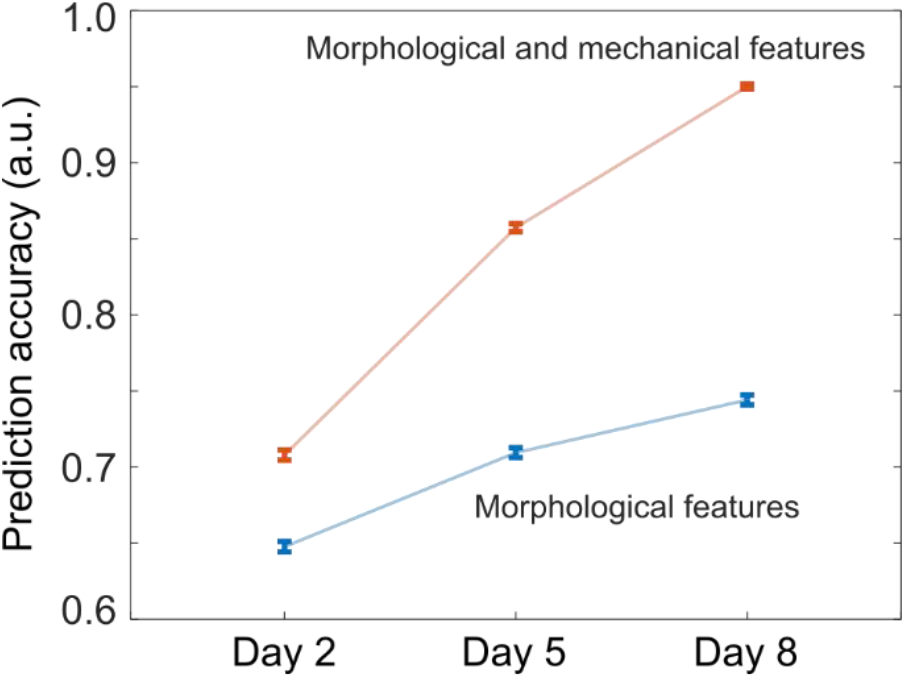
Prediction accuracy of cancer cell classification using a machine learning model. Error bar: standard error of the mean.

### 2.4 Deep-learning algorithm for cancer cell classification based on Brillouin images

#### 2.4.1. DenseNet121 models performance on the test set

We proceed to classify cancer and normal spheroids of each day using deep learning methods (see Experimental Section). During the validation phase, the F1-score corresponding to the cancer cell classification was used as the primary metric for performance evaluation, as detailed in Table 2. The results indicate a progressive enhancement in the F1-score, ascending from 0.645 to 0.970 with an increase in the temporal parameter. This trend may be attributed to the augmented differentiation of cellular structures of normal and cancerous spheroids over time. Specifically, the morphological characteristics of the normal cells exhibit an increasingly smooth periphery, coupled with a stabilization of pixel intensity within the cellular interior.

**Table 2.** Test results of spheroids using deep learning methods.

| Day of culture | Recall | Precision | F1-score |
| --- | --- | --- | --- |
| Day 2 | 0.625 | 0.667 | 0.645 |
| Day 5 | 0.882 | 0.750 | 0.811 |
| Day 8 | 1.000 | 0.948 | 0.970 |

#### 2.4.2 Explanation of the deep learning model prediction

We applied Gradient SHAP to explain the way our model makes predictions.^[48]^ The application of Gradient SHAP facilitates a granular understanding of the image regions that exert substantial influence over the model’s predictive accuracy. Gradient SHAP assigns a value ranging from 0 to 1 to each pixel in an image, indicating its importance in determining the output of the model. The higher this value, the more significant the pixel’s contribution to the model’s output. Illustrative results depicted in Figure 5 reveal a pronounced emphasis on pixel regions located within the cellular body. The heatmap corresponds to the foreground information (i.e., features) that the deep learning models used to make predictions. This observation substantiates the ability of our trio of models (pipeline shown in **Figure S3**, Supporting Information) to discern distinctive patterns corresponding to divergent temporal instances and to accurately categorize the input imagery. The explanation also validates that the reason for the increase in the F1-score is that the contour and the pixel intensity (i.e., Brillouin shift) within the cell become more distinctive, confirming the importance of the mechanical features extracted from Brillouin images.

**Figure 5.**
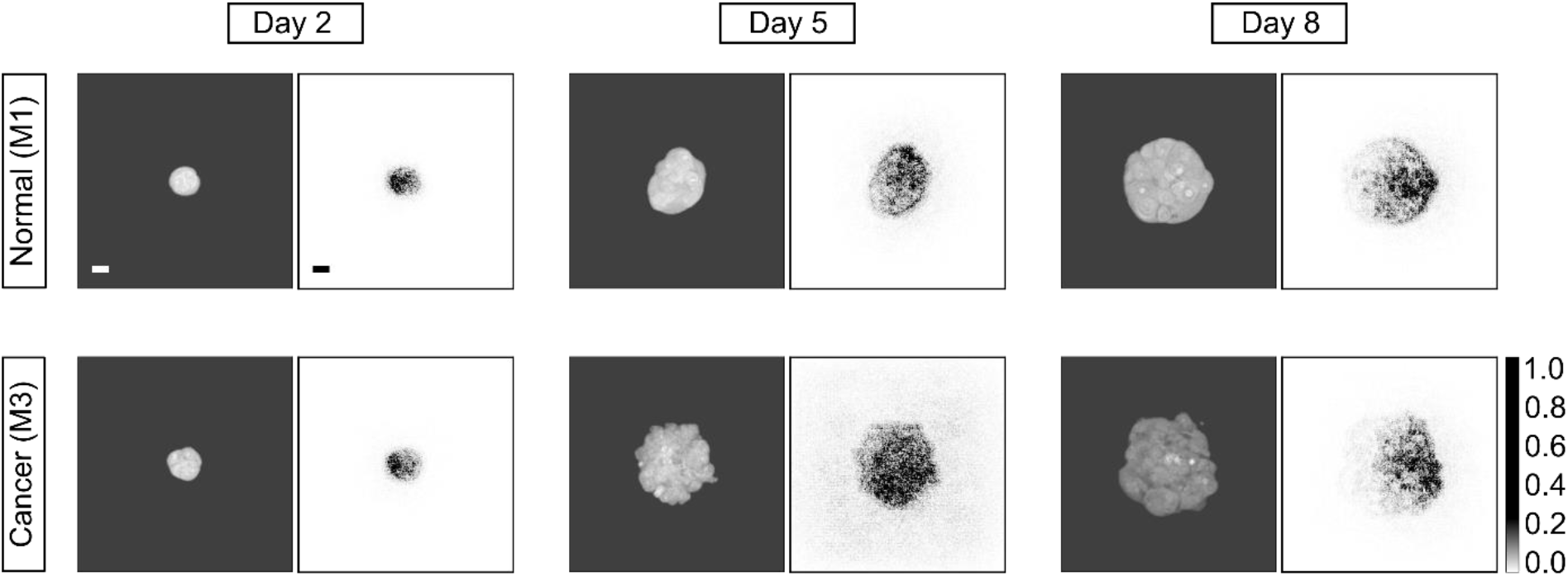
Representative results of interpreting outputs of the deep learning model using Gradient SHAP (SHapley Additive exPlanations). For each spheroid, the left panel is the normalized Brillouin image, and the right panel is the gradient map. Scale bar: 20 μm.

## 3. Discussion

The culture of MCF10A cells in 3D microenvironments leads to the formation of acini-like spheroids that closely mimic the glandular structure observed *in vivo*, as extensively documented in literature.^[42, 49]^ In alignment with prior findings, we captured the highly polarized arrangement of cells in normal spheroids from the acquired Brillouin images. The observed continuing increase in Brillouin shift of normal spheroids over an eight-day period is likely related to the increased cell number and/or intercellular compressive stress.^[39, 50]^ Intriguingly, metastatic cancerous spheroids exhibited distinct mechanical characteristics during the same culturing timeframe. Initially, from Day 2 to Day 5, the Brillouin shifts of cancerous spheroids rose at a rate comparable to that of normal spheroids, resulting in a minimal difference between cancer and normal spheroids. Given that single cancer cells also showed a lower Brillouin shift than normal cells on Day 0 (**Figure S2**), the observed differences between normal and cancerous spheroids during early-stage growth (i.e., Day 2 and 5) possibly arise from the individual cells. Conversely, a significant decrease in the Brillouin shift of cancerous spheroids was observed from Day 5 to Day 8, causing a widened difference in Brillouin shift between cancerous and normal spheroids on Day 8. Previous studies have shown that cancer cells exhibit a mesenchymal phenotype and reduced intercellular adhesion that facilitate metastatic spread. ^[10]^ Indeed, we observed that cancerous spheroids exhibited a larger average projection area than their normal counterparts and formed invasive peripheral protrusions by Day 8, which may reduce intercellular mechanical interaction. Therefore, it is likely that the mechanical variation observed between normal and cancerous spheroids on Day 8 is caused by both individual cells and altered intercellular interactions.

The dense and subcellular mechanical mapping from Brillouin microscopy enables us to extract spatially resolved mechanical features that were previously inaccessible. Here, we used the standard deviation and the skewness of the histogram distribution to quantify the heterogeneous biomechanics within the spheroid’s body. With a machine-learning based classification algorithm, we demonstrated that this additional information extracted from Brillouin images can substantially improve the accuracy of classifying cancerous spheroids. This finding was further confirmed by a deep learning pipeline that used solely Brillouin images for classification. Beyond cancer classification, we anticipate our results can be utilized to understand the underlying biomechanical mechanisms regulating tumor progression, such as the epithelial-to-mesenchymal transition,^[51, 52]^ the onset of metastasis,^[12, 53]^ and the formation of tumor,^[54, 55]^ in which mechanical interactions play crucial roles.

The direct readout from Brillouin microscopy is the Brillouin frequency shift, which is linked to the elastic longitudinal modulus through the material’s parameters, including density and refractive index. When the absolute value of modulus is needed, it is necessary to measure the Brillouin shift along with the refractive index *n* and/or density *ρ*. Nevertheless, for some biological cells and tissue, the refractive index and density changes in the same direction,^[31, 56, 57]^ resulting in a secondary variation in their ratio 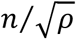. Therefore, the observed changes in Brillouin shift are primarily due to alternations in elastic longitudinal modulus. In this case, Brillouin shift has been validated as a good proxy for the relative change of mechanical modulus. ^[36, 58]^ Here, our work has demonstrated that Brillouin shift can serve as an effective contrast mechanism in label-free imaging. This paves the way for Brillouin microscopy to be a promising new imaging modality in cancer cell classification and detection.

## 4. Conclusion

We tracked the mechanical evolution of cancer and normal cells in a physiologically relevant 3D microenvironment by conducting longitudinal Brillouin imaging with growing spheroids. We found that cancerous spheroids exhibited distinct time-varying mechanical features, which are likely related to variations in cell behaviors and cell-cell interactions when compared to normal spheroids. Using an established machine learning algorithm and a customized deep learning pipeline, we demonstrate that these unique mechanical features hold promise for differentiating cancer cells from normal ones. Since force and mechanical modulus interact with each other and collectively regulate many physiological and pathological processes through mechanotransduction, mapping the spatial distribution of mechanical modulus can also provide insight into the generation and transmission of force, as well as the activation of signaling pathways. Given the important role of cell biomechanics in tumor progression, this work can help elucidate the biomechanical regulation in early-stage formation of tumors. Furthermore, it sheds light on using Brillouin-derived mechanical features as new biomarkers in cancer classification and detection.

## 5. Experimental Section

### Cell Culture

Non-tumorigenic MCF10A (M1) and malignant MCF10CA1h (M3) epithelial cells were generously provided by Barbara Ann Karmanos Cancer Institute (Detroit, MI, USA), and cultured according to protocols provided. All cells used did not exceed 10 passages. M1 cells were cultured in complete growth medium containing DMEM/F-12 (11330-032, Thermo Fisher Scientific), 5% horse serum (16050-122, Thermo Fisher Scientific), 20 ng/ml EGF (AF-100-15-1MG, Peprotech), 0.5 μg/ml hydrocortisone (H0888-1G, Sigma-Aldrich), 100 ng/ml cholera toxin (C8052-2MG, Sigma Aldrich), 10 μg/ml insulin (I1882-100MG, Sigma Aldrich), and 1% penicillin/streptomycin (15070-063, Thermo Fisher Scientific). M3 cells were cultured in minimal medium consisting of DMEM/F-12, 5% horse serum, and 1% penicillin/streptomycin. Both cell lines were cultured in T25 flasks at 37°C and 5% CO_2_. Both M1 and M3 spheroids were maintained in assay medium containing DMEM-F12, 2% horse serum, 5 ng/ml EGF, 0.5 ug/ml hydrocortisone, 100 ng/ml cholera toxin, 10 μg/ml insulin and 1% penicillin/streptomycin.

### Spheroids culture

The 3D on-top method was employed for spheroid colony formation, following established protocols.^[42]^ The Matrigel was thawed on ice overnight at 4°C. Ice cold pipette tips were used to ensure Matrigel was kept chilled. A volume of 200 μl of ice-cold Matrigel (354230, Corning) coated the 21 mm reservoir of a chilled 35 mm gridded dish (81166, Ibidi) and then placed in the incubator at 37°C and 5% CO_2_ for 30 minutes allowing the layer of Matrigel to solidify. Both MCF10A and MCF10CA1h cells were passaged individually at 75-80% confluency, with 0.05% trypsin: 0.53 mM EDTA (25-052-C1, Corning). Cells were then resuspended in resuspension medium, placed in a 15 ml conical tube, and centrifuged at 150 g for 5 minutes. A hemocytometer was used for cell counting, 0.5 × 10^4^ cells were suspended in 500 μl assay medium and carefully pipetted over the Matrigel layer and incubated for 30 minutes. Low seeding density was used for the 21 mm reservoir to avoid spheroids from clustering or growing in close proximity. An additional 500 μl of assay medium containing 10% Matrigel was added on top of the cell seeded Matrigel layer for a final concentration of 5% Matrigel. Cells begin to form acini-like spheroid colonies within a couple of days. Spheroid culture medium was changed the day before each experiment.

### Live/Dead Assay

To observe whether the 660 nm laser resulted in photodamage, we conducted a Live/Dead assay on day 9 using the same dishes used for Brillouin measurements. Spheroid medium was replaced with phenol red free DMEM assay medium (21063029, Thermo Fisher Scientific), prepared the same way mentioned above, excluding horse serum. Spheroids were stained with a 10 ml phenol red free assay medium mixture containing 1 μM ethidium bromide (EtBr, E1510, Sigma Aldrich), and 5 μM calcein AM (C3099, Invitrogen) and then placed in the incubator at 37°C and 5% CO_2_ for at least one hour. The spheroids were then washed and replaced with PBS. For nuclear staining, 1 drop/ml of Hoechst 33342 (R37605, Thermo Fisher) was added to each dish. Biological repeats of live/dead assays were conducted 3 times for spheroids measured with Brillouin as well as positive controls. To calibrate the fluorescence intensity of the red channel, we utilized dead spheroids as a negative control, which was prepared by replacing medium with -20ºC methanol for 20 minutes. During the live/dead experiments, spheroids were maintained at 37°C and 5% CO_2_. To quantify the fluorescence intensity of live cells (green fluorescence) and dead cells (red fluorescence), the corrected total cell fluorescence was calculated.

### Brillouin microscopy

Brillouin microscopy is based on the physical principle of spontaneous Brillouin light scattering, where the scattered light experiences a frequency shift (i.e., Brillouin shift) due to the interaction between the incident light and acoustic phonons generated by thermal fluctuation within a material. Brillouin shift *ω*_*B*_ is related to elastic longitudinal modulus *M*′ by

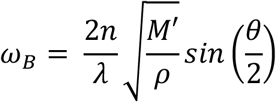

where *λ* is the wavelength of incident light, *n* and *ρ* are refractive index and density of the material, and *θ* is the scattered angle (*θ* = 180° for confocal Brillouin microscopy). The longitudinal modulus *M*′ is defined as the ratio of uniaxial stress and uniaxial strain.

A self-built confocal Brillouin microscope was used in this work. The detailed configuration of the Brillouin microscope has been reported before.^[35]^ In brief, a 660-nm continuous wave laser of 20-30 mW was used as light source. The laser beam was focused into the sample by an objective lens (40×/0.6, Olympus), yielding a spot size of 0.7×0.7×2.5 μm. The collected Brillouin signal was analyzed by a two-stage virtually imaged phased array (VIPA) spectrometer and recored by a EMCCD camera (iXon, Andor). During experiments, the Brillouin spectrometer was calibrated using standard materials (i.e., water and methanol). For spheroid and single-cell experiments, samples were placed into a on-stage incubator (UNO-T-H-CO2, Okolab) with 37ºC and 5% CO_2_. Brillouin images were acquired by scanning the sample with a step size of 0.5-2 μm and an acquisition time of 50 ms.

### Statistical Analysis

The unpaired two-tailed Welch’s t-test was used for comparison between cancer and normal cells under the same conditions. The paired two-tailed t-test was used for comparison between the same spheroids measured on different culturing days. For longitudinal Brillouin mapping of spheroids, at least 5 biological replicates were conducted for each cell line.

### Machine learning model

The support vector machine algorithm, provided by the open-source machine learning library Scikit-learn, was utilized for cell classification. First, geometric and mechanical features were extracted from Brillouin images, as outlined in **Table 1**. Next, we prepared two datasets for training the models: the first comprised solely geometric features, while the second included both geometric and mechanical features. Following established methodologies, we employed the Stratified K-Fold cross-validation method to split the dataset into 10 groups, where 9 groups were used for training the model, and 1 group for testing the model. After that, data was standardized by removing the mean and scaling to unit variance. Next, we conducted an exhaustive search to identify hyperparameters that maximized the performance of the model. Utilizing these optimal hyperparameters, the model was trained and tested on the training groups and testing group, respectively. To ensure the reliability of performance metrics, we trained 1000 models by conducting 100 runs with 10-fold rotations for each dataset. The performance of the model was assessed based on the average testing accuracy derived from all models.

### Deep-learning model

#### Model pipeline

Our model’s pipeline is shown in **Figure S3**. We trained three different DenseNet121 models for Brillouin data from Day 2, 5, and 8, which stands for Densely Connected Convolutional Network with 121 layers. In the process, as the Brillouin image propagates through the dense blocks, a rich feature hierarchy is built. The feature maps are sampled in transition layers. And after the last dense block, the global average pooling compresses the feature maps into a single vector that retains essential discriminative features, which is then used for classification. In the development of deep learning pipeline for the classification task of normal/cancer cellular structures, we optimized three distinct DenseNet121 architectures, obtained from the Medical Open Network for AI (MONAI). The optimization process employed Cross-Entropy Loss as the criterion for loss computation and utilized Adam optimization algorithm for the iterative refinement of model parameters. To mitigate the potential risk of overfitting, a series of data augmentation strategies, including random rotations (RandRotate) and random flips (RandFlip), were systematically employed to augment training set and integrate into the training regimen. The fine-tuning phase for the DenseNet121 models was conducted with an initial learning rate set at 5 × 10^−5^, extending across a total of 175 epochs.

#### Explanation on the deep learning model prediction

Upon completion of the training phase, the interpretability of the model’s decision-making process was elucidated using Gradient SHAP, ^[48]^ an eXplainable Artificial Intelligence (XAI) methodology. This technique synthesizes the principles of Shapley Additive Explanations and Integrated Gradients^[59, 60]^ to decipher the model’s predictive outputs. Gradient SHAP essentially combines these two methods and uses a linear explanation model, which is a weighted linear combination of gradients where the weights are the input features minus their baseline values. The idea is to approximate SHAP values by averaging the gradients multiplied by the input feature values over random samples of the input features, between a baseline (or reference) and the actual input.

## Acknowledgements

This work is supported in part by the National Institutes of Health (K25HD097288, R21HD112663), National Science Foundation (CBET-2339278), and the University Research Grant from Wayne State University.

## Data Availability Statement

All the data that support the finding of this work are available from the corresponding authors upon request.

