## Supplemental Materials for "Mechanical evolution of metastatic cancer cells in three-dimensional microenvironment"

**Supplementary Note 1: Single cells in Matrigel have lower Brillouin shifts than spheroids**

To understand biomechanics at single-cell level, Brillouin images of individual normal and cancer cells were acquired approximately 1 hour post-seeding in Matrigel. We found that the average Brillouin shift of cancer cells was lower than that of normal cells (**Figure S2**), which is consistent with previous studies showing cell stiffness is inversely related to metastatic potential. Notably, for both cell types, single cells had lower Brillouin shifts compared to their spheroid counterparts on Day 2. In addition, the standard deviations of Brillouin shifts showed no significant difference between normal and cancer cells, suggesting comparable levels of heterogeneity within each cell line. However, the distribution of Brillouin shift in cancer cells showed less asymmetry than in normal cells, and single cells of both cell types exhibited less asymmetry in distribution compared to their spheroids on Day 2. In summary, the mechanical distinctions between normal and cancer cells persist from single-cell stage to multicellular spheroids by Day 2. Nevertheless, single cells had lower Brillouin shift and asymmetry in distribution compared to the spheroids.

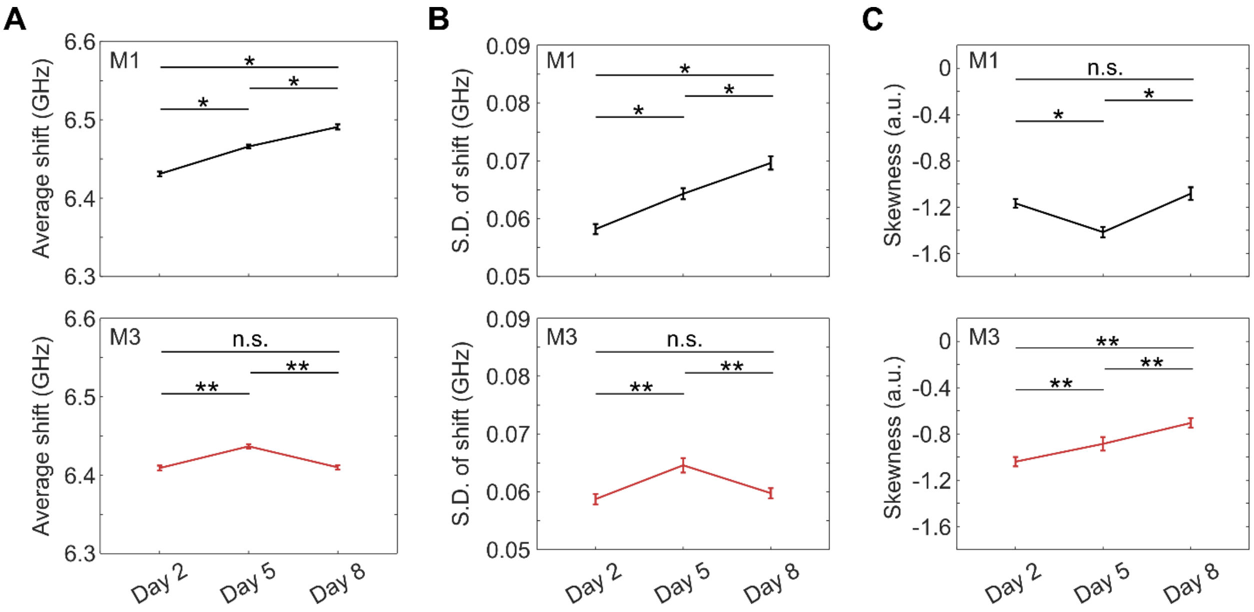


**Figure S1**. Time evolving mechanical features of normal (M1) and cancer (M3) spheroids. (A) Average Brillouin shifts. * $p=1.8\times10^{-17}$ (Day 2- Day5); $p=4.6\times10^{-14}$ (Day 5- Day8); $p=3.5\times10^{-29}$ (Day 2- Day8). ** $p=4.3\times10^{-15}$ (Day 2- Day5); $p=2.3\times10^{-15}$ (Day 5- Day8). (B) Standard deviation of Brillouin shifts. * $p=3.4\times10^{-9}$ (Day 2- Day5); $p=5.1\times10^{-5}$ (Day 5- Day8); $p=1.7\times10^{-13}$ (Day 2- Day8). ** $p=7.6\times10^{-5}$ (Day 2- Day5); $p=1.4\times10^{-4}$ (Day 5- Day8). (C) Skewness of distribution in Brillouin shifts. Error bar represents the standard error of the mean. * $p=9.4\times10^{-7}$ (Day 2- Day5);

*p*=4.9×$10^{-9}$ (Day 5- Day8). ** *p*=0.009 (Day 2- Day5); *p*=8×$10^{-4}$ (Day 5- Day8); *p*=2.1×$10^{-9}$ (Day 2- Day8). n.s.: no significant difference.

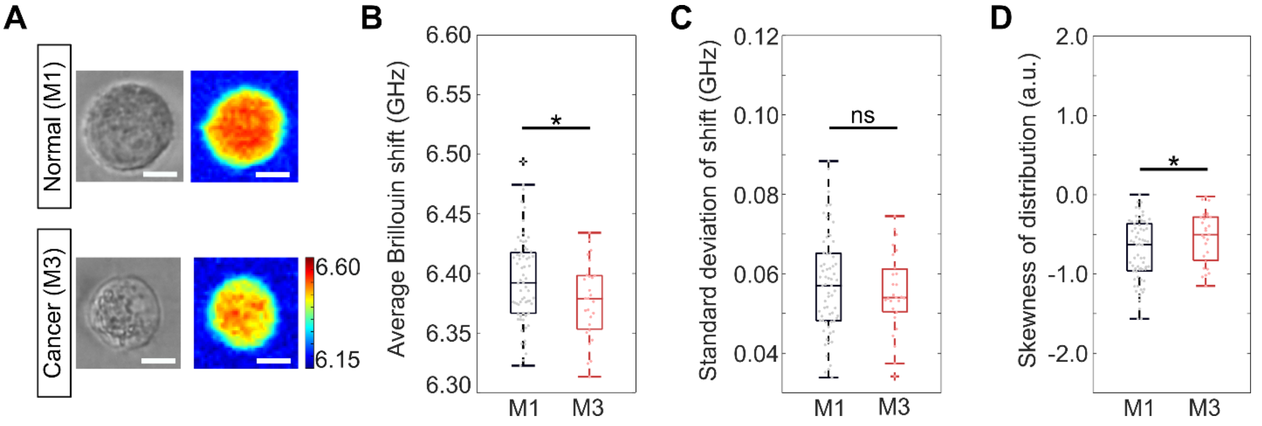


**Figure S2**. Single cell results. (A) Representative Brillouin and brightfield images of a normal cell (M1) and a cancer cell (M3) seeded in Matrigel. Scale bar: 10μm. (B) Average Brillouin shifts of normal (n=66) and cancer cells (n=27). * *p* = 0.0292. (C) Standard deviation of Brillouin shifts. ns: no significant difference. (D) Skewness of distribution in Brillouin shifts. * *p* = 0.0446.

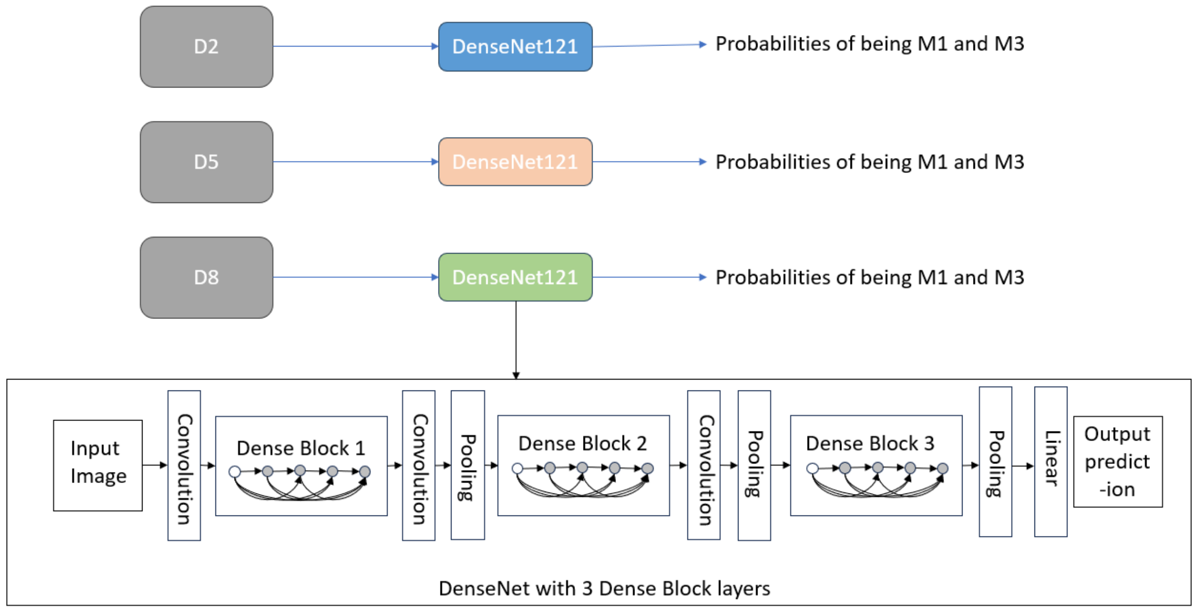


**Figure S3**. An illustration of the deep learning pipeline.
